# Determination of Absolute Intramolecular Distances in Proteins by Anomalous X-ray Scattering Interferometry

**DOI:** 10.1101/2024.02.09.579681

**Authors:** Samuel Stubhan, Anna V. Baptist, Caroline Körösy, Alessandra Narducci, Gustavo Gabriel Moya Muñoz, Nicolas Wendler, Aidin Lak, Michael Sztucki, Thorben Cordes, Jan Lipfert

## Abstract

Biomolecular structures are typically determined using frozen or crystalline samples. Measurement of intramolecular distances in solution can provide additional insights into conformational heterogeneity and dynamics of biological macromolecules and their complexes. The established molecular ruler techniques used for this (NMR, FRET, and EPR) are, however, limited in their dynamic range and require model assumptions to determine absolute distance (distributions). Here, we introduce anomalous X-ray scattering interferometry (AXSI) for intramolecular distance measurements in proteins, which are labeled at two sites with small gold nanoparticles of 0.7 nm radius. We apply AXSI to two different cysteine-variants of maltose binding protein in the presence and absence of its ligand maltose and find distances in quantitative agreement with single-molecule FRET experiments. Our study shows that AXSI enables determination of absolute intramolecular distance distributions under virtually arbitrary solution conditions and we anticipate its broad use to characterize protein conformational ensembles and dynamics.

## INTRODUCTION

Atomic resolution biomolecular structures are typically determined using frozen or crystalline samples with cryo-EM (1,2) and X-ray methods (3–6), respectively, or in aqueous solution at room temperature with NMR (7,8). Measurement of intramolecular distances can provide additional insights into the structure and dynamics of biological macromolecules and their complexes. A sufficient number of intramolecular distances enables the determination of high-resolution structures (9–13) and can also provide critical information about the conformational ensemble and dynamics of macromolecules (14–20) based on molecular ruler techniques such as PELDOR/DEER (EPR) (21–23) or single-molecule Förster resonance energy transfer (smFRET) (24–27).

Also small-angle X-ray scattering can provide information about intramolecular distances in biomolecules. Notably the *P(r)* distribution, i.e. the Fourier transform of the scattering intensity profile *I(q)*, is a histogram of pairwise distances and can be readily obtained from scattering data (28–30). However, *P(r)* does not contain information about which specific pair contributed to a given distance. Labeling macromolecules with electron-rich labels at two positions – e.g. heavy atoms (31–33), ions, or small gold nanocrystals – combined with SAXS as readout can overcome this limitation. In conventional X-ray scattering interferometry (XSI) (34–40) with gold labels, the label-label interference term is isolated from other scattering contributions by measuring multiple samples, including the double-labeled, two single-labeled, and unlabeled macromolecule (34,35). An alternative approach to separating the gold-gold term, termed anomalous XSI or AXSI, uses anomalous small-angle X-ray scattering (ASAXS) and relies on the energy-dependence of the gold scattering signal (32,33,41–43). A regularized Fourier transform of the gold-gold scattering term then directly provides the distribution of distances *P(d)* between the gold labels. (A)XSI has several advantages compared to other molecular ruler techniques: (i) It provides distance distributions on an absolute length scale, based on the fact that it is straight-forward to measure the momentum transfer *q* (*q =* 4 π sin(*θ*)/*λ*, where *2θ* is the total scattering angle and *λ* the X-ray wavelength) on an absolute scale; (ii) (A)XSI provides the full distribution of intramolecular distances (not only mean inter-label distances), without broadening through e.g., photophysics, as seen in FRET; (iii) it can readily be applied to distances > 10 nm, which remains very challenging for NMR, EPR (21,22,44), or FRET (45); (iv) finally (A)XSI distance measurements are not sensitive to label orientation or the specific label environment, unlike FRET approaches (17). Insensitivity of the distance measurement to the environment is advantageous for measurements to determine conformational changes in response to e.g., denaturant (46), salt (47–49), or ligand concentration. ASAXS-based AXSI measurements have the advantage that they only require preparation of the double-labeled sample, as opposed to traditional XSI, which requires matching single-labeled constructs as well. However, so far AXSI has only been established experimentally for DNA constructs, which can be labeled in a straightforward way (43).

Here, we demonstrate AXSI intramolecular distance measurements in proteins that undergo conformational changes upon ligand binding. We use MalE, the soluble periplasmic component of the maltose import system of *E. coli* (50–52), which has been characterized in detail previously by smFRET experiments and other structural methods (17,50–57). MalE undergoes a conformational change from an open/apo to a closed/holo state upon binding maltose with a dissociation constant K_d_ of ∼1-2 µM (17). We analyze AXSI data for two double-cysteine variants of MalE and extract distance distributions of the apo and holo states that exhibit sharp main peaks. The main peak position can be determined with Ångström precision and the measured distances are in good agreement, within experimental error, with quantitative distance determination via smFRET.

## RESULTS AND DISCUSSION

### Labeling of MalE mutants via thiol-gold chemistry

(A)XSI measurements for computing intramolecular distances necessitate site-specific attachment of gold-labels. For this purpose, we used variants of the 42.4 kDa maltose-binding protein MalE that comprise two cysteines in the two different lobes of the protein (Supplementary Materials and Methods), either at positions 31 and 212 (MalE_31-212_; Figure 1) or at positions 36 and 352 (MalE_36-352_) (17,52). MalE lacks native cysteines, thus allowing site-specific attachment of gold labels via chemical coupling to thiols (52) (Figure 1a). Thioglucose coated gold nanoparticles (NPs) were synthesized following a one-phase Brust-Schiffrin method (58) (Supplementary Materials and Methods) and exhibit a radius of 0.7 nm with high monodispersity (34,35,40) (Supplementary Figure S1). Proteins were labeled with the NP on nickel-sepharose columns (23,52,59) and subsequently purified by size exclusion chromatography (Supplementary Figures S2 and S3).

**Figure 1.**
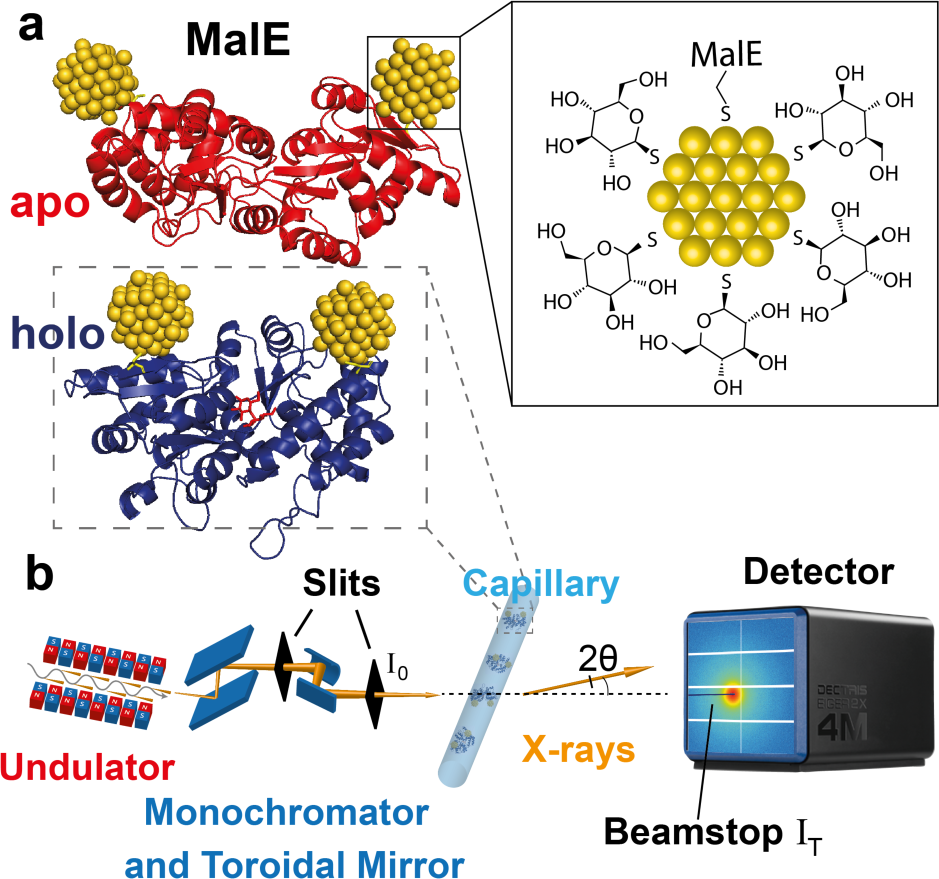
Schematic of anomalous X-ray scattering measurements to determine intramolecular distance distributions. **a)** Illustrations of double-labeled MalE in the apo and holo state with gold labels at amino acid positions 36 and 352 (rendered from PDB ID 1OMP (red – apo) and 1ANF (blue – holo), respectively. Gold nanocrystals are positioned using FPS calculations (12)). The zoom depicts the thioglucose shell on the gold NPs as well as the S-Au attachment to the protein. **b)** Illustration of the SAXS experiment. The undulator and X-ray optics at the synchrotron beam line provide X-rays with tunable energy. The monochromator is used to select energies and collimated X-rays being are scattered by the sample in a quartz capillary. The incident intensity I_0_ is measured in front of the capillary and the transmitted intensity I_T_ is measured at the beamstop. Scattered photons are collected in an Eiger2X 4M pixel detector.

### SAXS and ASAXS measurements of MalE constructs

We carried out SAXS measurements at both fixed and variable X-ray energies at beamline ID02 of the European Synchrotron Radiation Facility (ESRF; (60–63) and Supplementary Materials and Methods; Figure 1). Control measurements of the unlabeled protein at fixed energy reveal SAXS profiles indicative of a monodisperse sample and show systematic, but subtle changes upon addition of 10 mM maltose, with radii of gyration in good agreement with predictions from the crystal structures in the open and closed conformations (Supplementary Figure S4).

ASAXS data were recorded for double-labeled, single-labeled, and unlabeled MalE constructs by recording scattering profiles at 9 energies around the gold L-III absorption edge at 11.919 keV (Figure 2). For all ASAXS measurements, ascorbic acid was added to the buffer to reduce radiation damage (Supplementary Figure S5). The scattering profiles at different energies for the double-labeled MalE_31-212_ and MalE_36-352_ constructs both show oscillations (Figure 2a), in particular in the range *q =* 0.1-0.2 Å^−1^, which are absent in the unlabeled data (Supplementary Figure S4), indicative of the gold-gold interference contribution (40). The oscillations shift upon addition of maltose (Figure 2a, inset), suggesting a modulation of gold-gold interference term upon addition of maltose. Further, the scattering profiles show systematic changes with X-ray energy: the intensity decreases when approaching the L-III absorption edge.

**Figure 2.**
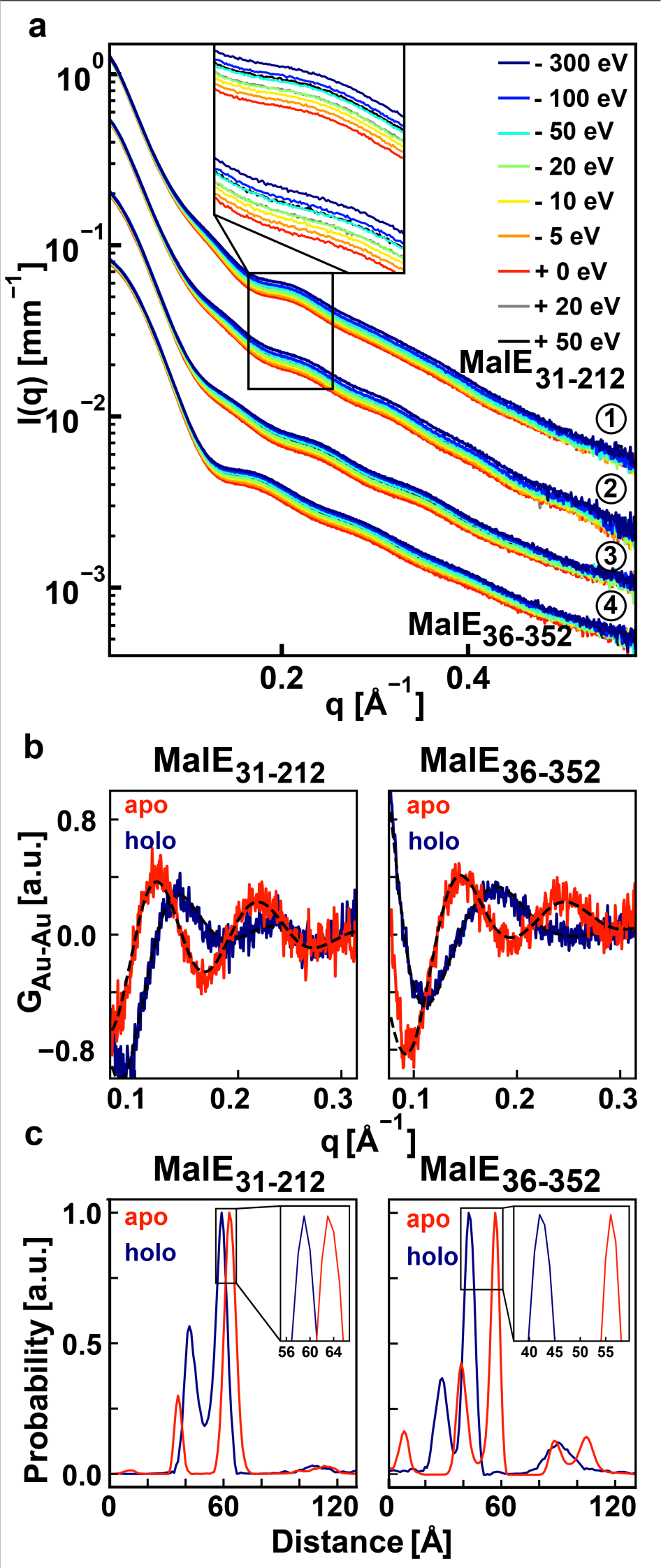
ASAXS data and distance distribution for MalE labeled at position 36 and 352 and MalE labeled at position 31 and 212. **a)** SAXS measurements of both double-labeled MalE mutants with and without maltose measured at 9 energies around the gold L-III absorption edge. ① MalE_31-212_ ② MalE_31-212_ with 10 mM maltose ③ MalE_36-352_ and ④ MalE_36-352_ with 10 mM maltose. Data are vertically offset for clarity (by scaling factors ①:10, ②:6, ③:2 and ④:1). Indicated energies are relative to the Au L-III edge. **b)** Gold-gold scattering interference terms for MalE_36-352_ (left) and MalE_31-212_ (right) in the absence and presence of 10 mM maltose after correction for 60% single-labeled and 30% unlabeled contributions. **c)** Distance distributions *P*(d) obtained by maximum entropy inversion from the interference terms in panel b. The insets show a zoom on the main peaks in the distance distributions.

HPLC purification of the double-labeled sample removes dimers and aggregates from the monomer peak, however, single-labeled and unlabeled species remain in the solution, visible as bands in gel electrophoretic analysis of the sample. From the gel, we estimate ∼60 % unlabeled, ∼30 %, single-labeled, and ∼10% double-labeled sample, which agrees with a protein:gold NP concentration ratio of 1:0.5 (Supplementary Figure S2 and S3). Despite of the relatively low labeling efficiency, we still get a robust signal, since the gold particles scatter strongly with ∼85 atoms and a ∼40 times higher electron density contrast compared to proteins. Additionally, in principle, only the double-labeled sample will contribute a gold-gold term to the scattering pattern. However, the presence of unlabeled and single-labeled species might deteriorate the signal. Since we conducted ASAXS measurements also for single-labeled mutants MalE_31_, MalE_36_, MalE_212_, MalE_352_ and unlabeled samples, we can subtract their scattering contributions from the double-labeled sample (Supplementary Figure S6). We tested and refined the influence of subtracting single- and unlabeled protein contributions in the ASAXS data analysis (see below).

### Determination of the gold label-gold label distance distribution from ASAXS data

We analyzed the ASAXS data determined the gold-gold scattering contribution from the corrected and energy-dependent scattering data with a matrix inversion approach described previously (42,43). The approach takes into account the atomic scattering factor of gold and the form factor of the gold spheres (Supplementary Figure S1) and exploits the fact that the atomic scattering factors for non-gold atoms show minimal energy dependence within the chosen energy range (Supplementary Methods and Supplementary Figure S7). The matrix inversion yields the gold-gold structure factor *G_Au-Au_*, which was corrected by a constant offset by subtracting the mean (43) (Figure 2b) and Fourier transformed with a maximum entropy algorithm (34,35,40,43) to obtain the real-space distance distributions *P*(d) (Figure 2c).

Using this procedure, we determined the gold label-gold label distance distributions *P*(d) for the two MalE variants, MalE_31-212_ and MalE_36-352_, both in absence and presence of a high (10 mM) concentration of maltose (Figure 2c). Under all conditions, the *P*(d) distributions exhibit a major peak and additional, smaller peaks at smaller and larger distances. We find that peaks at smaller and larger distances (> 80 Å) are variable from data set to data set and are sensitive to details of the single- and unlabeled subtraction and maximum entropy procedure and are likely due to imperfections of the experimental data (40). In contrast, the positions of the main peak in either condition are robust (Supplementary Table S1, S2 and Supplementary Figure S8).

### Analysis of uncertainty in AXSI measurements

To determine the uncertainties of the measured distance distributions, we analyzed the variability introduced by several factors. First, we tested the uncertainty introduced by the maximum entropy algorithm used to compute the *P(d)* distributions. For each measured gold-gold term *G_Au-Au_*, we carried out 20 repeat runs of the maximum entropy algorithm and fitted the main peak with a Gaussian to determine its position and standard deviation (Supplementary Figure S9). We find only small deviations between repeat runs of the maximum entropy algorithm, with standard deviations of the main peak positions of 0.5 Å on average.

Next, we test the sensitivity to subtracting unlabeled and single-labeled contributions to the scattering pattern. We subtracted varying quantities of single-labeled and unlabeled protein contributions from the scattering pattern and computed *P*(d) functions as described above. We find that the secondary peaks in the distance distribution are smallest if 60 percent unlabeled and 30 single-labeled contributions are subtracted (Supplementary Figure S8 and data shown in Figure 2) for both mutants MalE_36-352_ and MalE_31-212_, which agrees with the estimated fractions from gel analysis (Supplementary Figure S2). Importantly, adjusting the amount of single and unlabeled contributions subtracted or even using the scattering data without subtraction affects only the level and position of the secondary peaks and changes the main peak position by at most 1 Å (Supplementary Table S1, S2). Thus, our analysis suggests that while the unlabeled and single-labeled contributions add to the level of experimental noise and affect the exact shape of the *P*(d) function, the main peak corresponding to the gold label-gold label distance is robust against these perturbations. Finally, we tested the reproducibility of the AXSI measurement by performing a repeat measurement for each sample. We find that the mean peak position for repeat measurements only varies by at most 0.7 Å (Supplementary Table S1, S2). Taken together, our error analysis suggests that we can determine the center of the main peak in the *P*(d) distributions to better than 1 Å (Table 1 and Supplementary Tables S1 and S2), in agreement with previous analyses of (A)XSI measurements for nucleic acids (34,40,43). As a control, we determined the gold-gold structure factor *G_Au-Au_* using conventional XSI (34,35,40), i.e. with measurements at only a single X-ray energy of the double-labeled, both single-labeled, and the unlabeled protein samples with SAXS. The control measurement for MalE_36-352_ in the absence of maltose gives a main peak in the distance distributions in excellent agreement with the results of the AXSI analysis (Supplementary Figure S10).

**Table 1.** Intramolecular distances in MalE_31-212_ and MalE_36-352_ in the absence (apo state) and presence of 10 mM maltose (holo state). Values reported for AXSI are the mean ± standard deviation of the peak position averaged over repeat measurements and different levels of background subtractions (Supplementary Tables S1 and S2). Values for FRET are the mean ± standard deviation from three technical repeats taking into account the uncertainties of Förster radius (assuming a Förster radius of 5.1 nm (23)) and quantum yield. Simulated values are the mean label distances ± the standard deviations of simulated distances. For FRET labels we report the results of the established procedures FPS and FRETraj; the coarse-grained simulations are very similar to the FPS results. For AXSI labels we conversely report values from FPS and the coarse-grained method, since FRETraj gave very similar values to FPS (Figure 4 and Figure S12).

|  | MalE<sub>31-212</sub><br>apo | MalE<sub>31-212</sub><br>holo | MalE<sub>36-352</sub><br>apo | MalE<sub>36-352</sub><br>holo |
| --- | --- | --- | --- | --- |
| **Measurements (mean ± error) [Å]** |  |  |  |  |
| AXSI | 63.7 ± 0.5 | 59.4 ± 0.5 | 56.0 ± 0.3 | 43.1 ± 0.6 |
| FRET | 63.0 ± 2.4 | 56.3 ± 2.2 | 54.1 ± 2.1 | 44.9 ± 1.7 |
| **Simulations (mean ± standard deviation) [Å]** |  |  |  |  |
| FPS AXSI | 67.9 ± 0.5 | 59.7 ± 0.4 | 60.7 ± 1.2 | 44.6 ± 0.7 |
| Coarse-grained AXSI | 64.4 ± 2.8 | 56.6 ± 2.0 | 59.0 ± 4.4 | 42.7 ± 4.5 |
| FPS FRET | 68.6 ± 8.3 | 56.7 ± 9.5 | 60.5 ± 11.3 | 44.3 ± 11.3 |
| FRETraj FRET | 63.0 ± 9.6 | 53.8 ± 10.0 | 56.8 ± 11.6 | 42.2 ± 11.8 |
| C<sub>β</sub> distance | 51.5 | 42.6 | 50.6 | 39.8 |

### Determination of MalE intramolecular distances by smFRET

To provide a reference and enable direct comparison of the results with an established technique, we performed distance measurements on MalE_31-212_ and MalE_36-352_ by single-molecule FRET using alternating laser excitation (ALEX) (64–66) (Figure 3a and Supplementary Methods). We employed a data analysis approach similar to a recent multi-lab FRET comparison study (17) to extract mean interprobe distances between donor (Alexa 555) and acceptor fluorophore (Alexa 647) from intensity-based single-molecule measurements (13,15,64–68) based on the Förster relation (Figure 3b). Both MalE_31-212_ and MalE_36-352_ are designed to show changes from larger to smaller interprobe distances upon maltose binding. We observe shifts from low to intermediate FRET efficiency values *E* from the apo to holo state, as expected (Figure 3c, d). The mean *E* values change from 0.22 to 0.36 for MalE_31-212_ and from 0.4 to 0.67 for MalE_36-352_ (Supplementary Figure S11), corresponding to a reduction in the calculated interprobe distances (Table 1).

**Figure 3.**
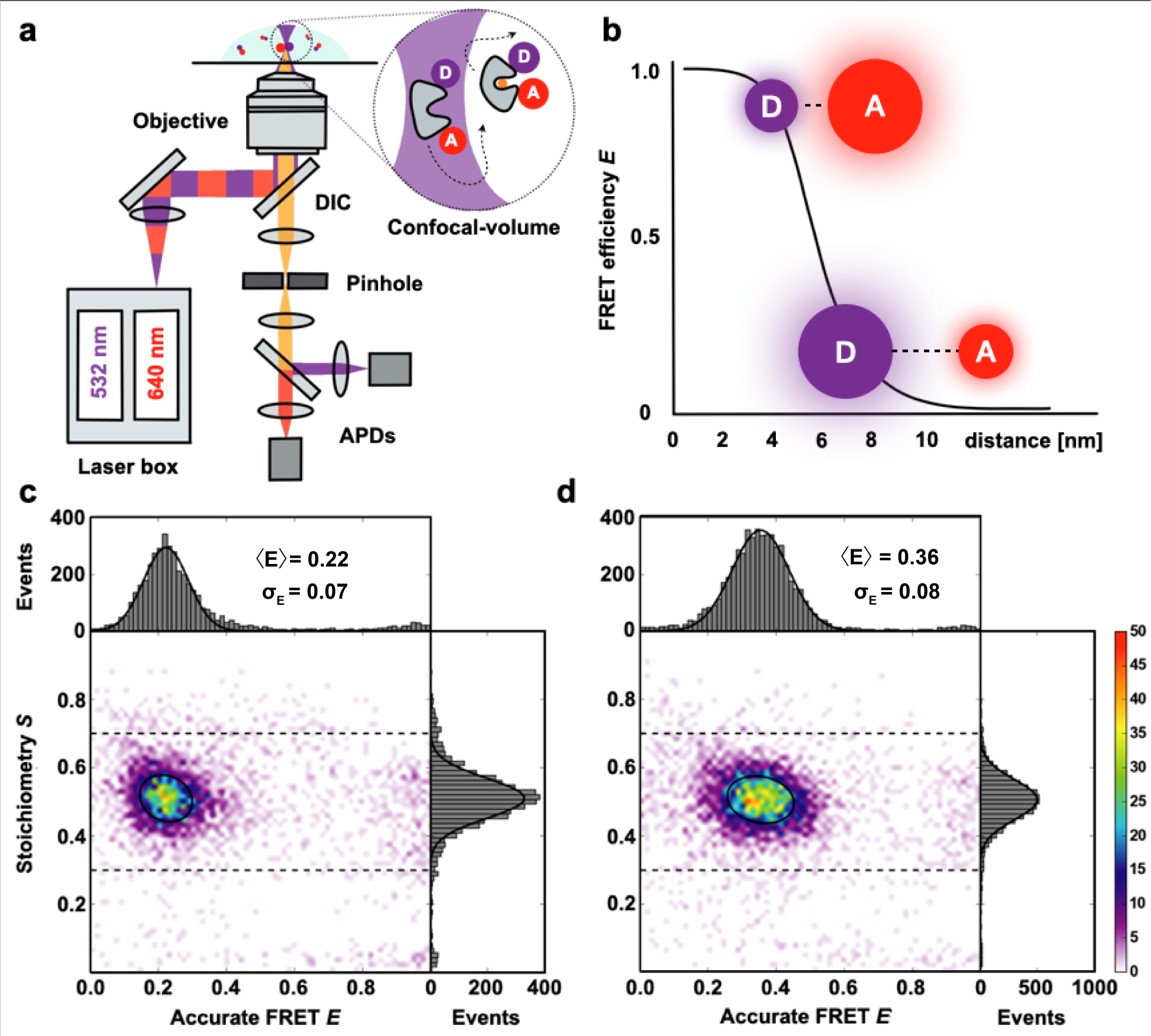
Monitoring conformational changes in MalE by smFRET with ALEX. **a)** Schematic overview of an ALEX confocal microscopy setup with two fiber-coupled modulated laser sources for alternating excitation of donor **(D)** and acceptor **(A)** dyes. The laser is expanded, collimated, and directed into an objective with a high numerical aperture through a dichroic mirror (DIC). The objective is used for excitation and detection of the fluorescence of individual proteins in a diffraction limited excitation spot. Subsequently, a pinhole spatially filters the fluorescence before it undergoes spectral separation into green and red detection channels. **b)** Schematic plot of the FRET efficiency (*E*) as a function of the distance between a donor fluorophore and an acceptor. **c, d)** Corrected accurate *ES*-histograms of MalE_31-212_ labeled with Alexa555 and Alexa647 depicting the ligand-free open (c) and the liganded closed (in the presence of 1 mM maltose; d) conformation.

### Modeling the label geometry for FRET and AXSI measurements

AXSI and FRET give mean distances that are in good agreement (Table 1). However, we note that both AXSI and FRET measure the distances between the respective labels and not directly between the positions of the labeled residues. Therefore, the attachment, size, and flexibility of the organic dyes or gold labels need to be considered when interpreting distance measurements (13,69). Taking into account the label geometries is particularly relevant in light of the very high resolution of AXSI measurements, where distances are determined to better than 1 Å (Table 1), which is much smaller than the label and linker sizes and also smaller than the spatial resolution of FRET which is on the order of 2-5 Å (17,70). Taking the crystallographic structures of MalE as a starting point, we simulate label positions using label parameters summarized in Supplementary Table S3 and following several different approaches (Figure 4). First, we calculated the label distances with FPS (“FRET positioning and screening” (12,71), which generates accessible volumes for each dye and computes the distance distribution assuming random sampling of the accessible volume (Table 1, “FPS”). Second, we use FRETraj (72) to compute distances based on accessible contact volumes (ACVs) (Table 1, “FRETraj”), which have been shown to provide a better estimate of label-label distances than the full accessible volume if the dyes interact with the protein surface (17). Finally, we compute accessible volumes using a simple coarse-grained sampling (see Supplementary Methods: Coarse-grained simulation of accessible volumes) and calculate the mean and standard deviation of distances in the sampled positions (Table 1 and Figure 4, Supplementary Figure S12).

**Figure 4.**
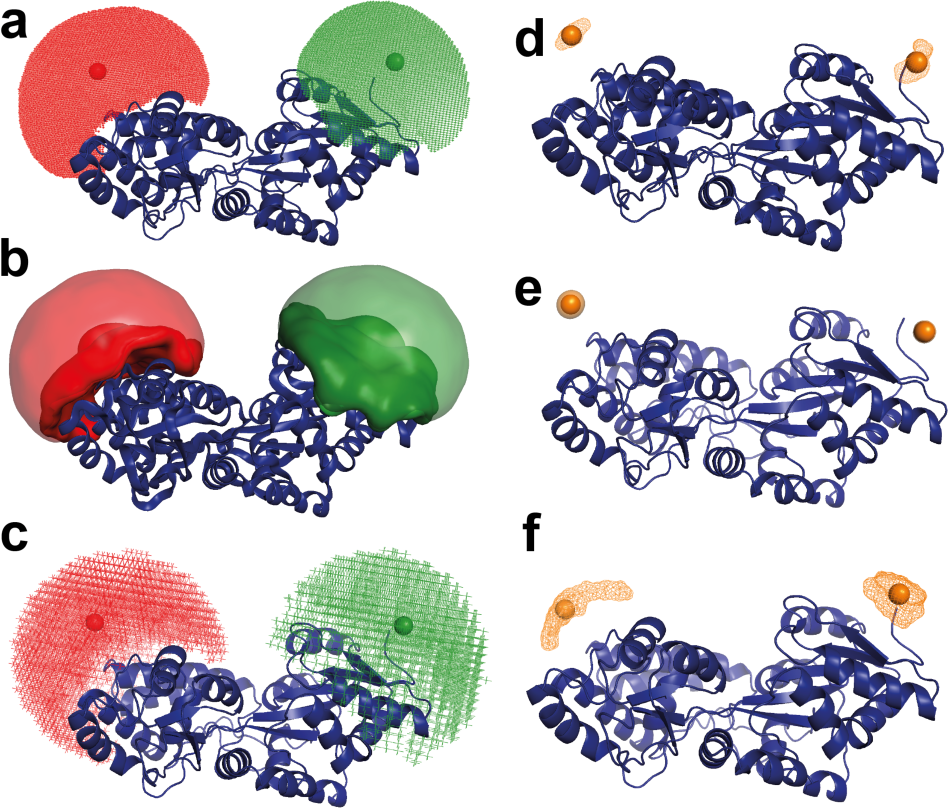
Simulations of label positions for FRET dyes and gold nanoparticles using FPS. (**12**)**, FRETraj** (**72**)**, and a coarse-grained computation of accessible volumes.** All panels show MalE_36-352_ in the apo state (PDB ID: 1OMP). **a)-c)** Positions for the FRET dyes Alexa555 and Alexa647 are shown as colored clouds as computed with FPS (a), FRETraj (b), and coarse-grained simulations (c). **d)-f)** Positions of the gold nanoparticles used for AXSI measurements computed with FPS (d), FRETraj (e), and coarse-grained simulations (f). The geometrical parameters used in the calculations are in Table S3. The resulting mean label-label distances and their standard deviations are in Table 1.

To model the label positions for AXSI measurements, we adopted the same procedures considering the size and attachment of the gold nanoparticles. The gold nanoparticles, in contrast to fluorescent dyes, are directly attached to the sulfur atom of the cysteine residues. Therefore, the C_β_–Au distance is only ∼3 Å. The attached gold NP have a radius of 7 Å, resulting in a distance from attachment atom C_β_ to the center of the label of ∼10 Å. The much more confined attachment of the gold nanoparticles compared to the fluorescent dyes used for FRET results in much more narrow predicted distance distributions: the predicted distributions for FRET labels have standard deviations of ∼10 Å in contrast to only ∼0.5-4 Å for our AXSI labels (Table 1).

### Comparison of distances from AXSI and FRET and modeling

The intramolecular distances determined experimentally by AXSI and FRET agree for MalE_31-212_ and MalE_36-352_ in both the apo and holo state within error (Table 1 and Figure 5). This close agreement, despite the different labels used, suggests that the different physico-chemical properties and their differences in geometry do not significantly affect the mean positions of the labels. Comparing the experimentally determined distances to the modeled distance distributions, we find good agreement of the mean distances in the closed or holo state for both MalE_31-212_ and MalE_36-352_. In the closed state, all of the approaches to model the mean distances give fairly similar results, in particular for MalE_36-352_, due to the direction of the label attachment being approximately perpendicular to the vector connecting the label centers (Figure 1a). In contrast, in the open or apo state the predicted distances tend to be larger than the experimentally determined values, in particular for MalE_36-352_ (Figure 5). We note that for the open conformation the details of the modeling play a larger role, compared to the closed conformation. In particular, the predictions based on the accessible surface volume using FRETraj fit the experimental data for FRET better than the accessible volume-based predictions, suggesting that the fluorescent labels might have some tendency to stick to the protein’s surface (17,71). The differences in the apo state might also indicate that the protein conformations in solution could deviate from the crystal structure. As a plausibility test, we used a simple elastic network model approach (73) to deform the structure of the open conformation of MalE (Supplementary Figure S13) and find that a simple deformation of the protein along the first normal mode could explain the observed difference between measured and predicted distances.

**Figure 5.**
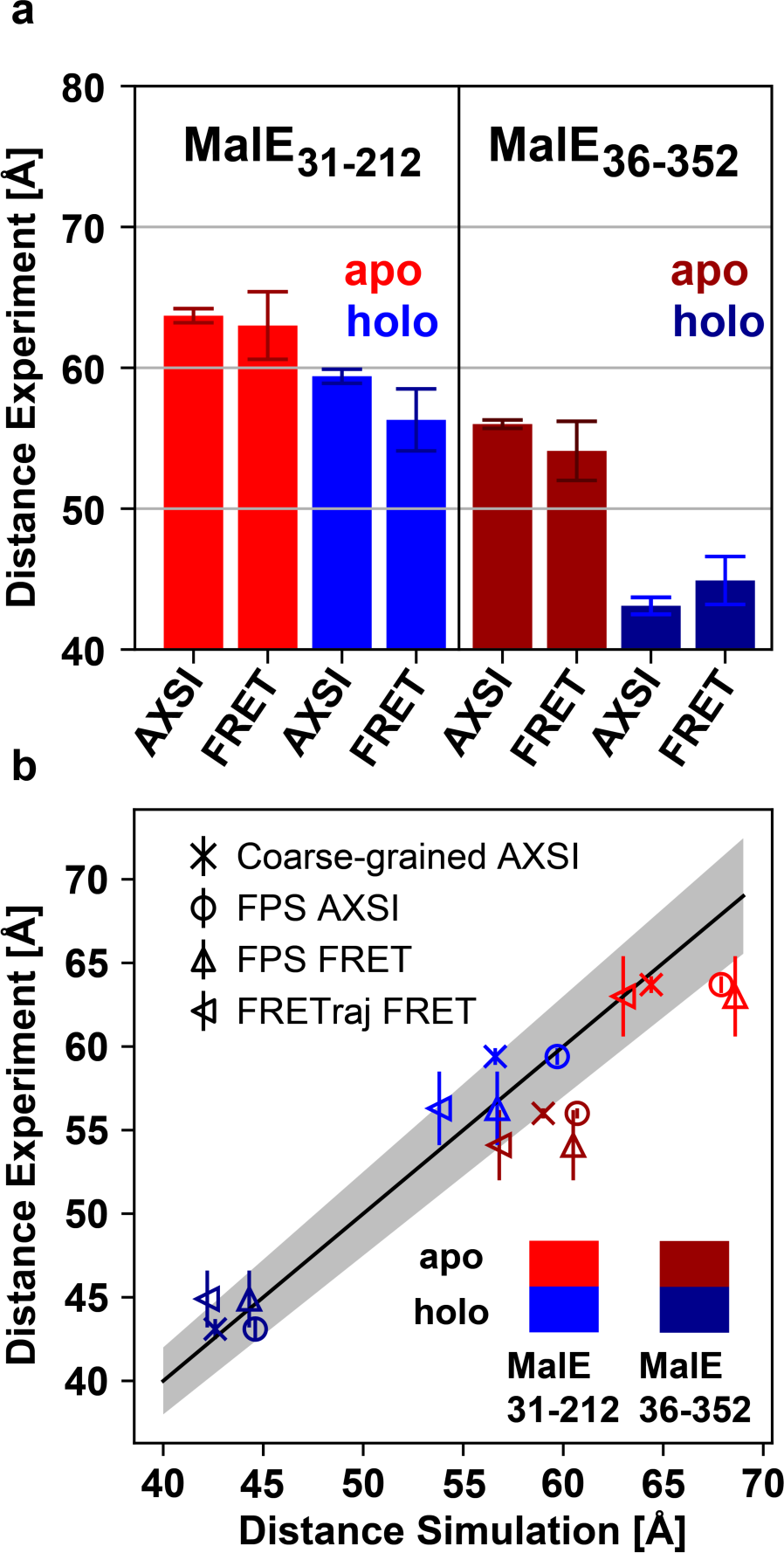
Comparison of AXSI and FRET measurements and structural modeling. **a)** Experimentally determined distances from AXSI and FRET for both MalE variants. Errorbars depict experimental errors (see main text). **b)** Comparison of experimentally determined distances and the structural models: coarse-grained for AXSI labels (crosses), FPS for AXSI labels (circles), FPS for FRET labels (upward triangles) and FRETraj for FRET labels (left triangles). Experimental uncertainties of the respective techniques are shown. The solid line marks a 1:1 relation and the grey area indicates 5% deviation.

Unlike in FRET measurements, AXSI provides, in principle, the full distance distribution (34,43). The mean peaks of the *P*(d) distributions are fairly narrow, with a width of typically ∼2 Å (Supplementary Tables S1 and S2), suggesting that the combined protein and label movement has a magnitude of ∼2 Å (except for MalE_36-352_ holo with ∼5 Å), which is similar to the distributions obtained by modeling the relatively inflexibly attached gold nanoparticles, assuming otherwise static protein structures. Our findings show that the labeled residues of MalE are rigid and adopt a fairly static conformation in solution, as might be expected for a well-folded, globular protein.

## CONCLUSION

In summary, we demonstrate accurate intramolecular distance measurements using AXSI for a protein that undergoes ligand-induced conformational motion. Mean distances can be determined very precisely (within < 1 Å) and we find excellent agreement with distances measured experimentally by quantitative FRET. In the future, improved labeling and purification procedures and more sophisticated modeling approaches, e.g. based on molecular dynamics simulations, should enable improved estimates of the full distance distributions and allow for full quantitative comparisons.

In conclusion, the introduction of AXSI for proteins opens up exciting possibilities in structural biology and beyond. The ability to determine intramolecular distance distributions for proteins in free solution and under virtually arbitrary solution conditions makes our approach ideally suited to resolve questions regarding partially folded conformations and natively disordered proteins (46,74–76). Beyond intramolecular distance measurements, AXSI might be extended to studying protein-protein interactions, ligand binding kinetics, and signaling pathways. Furthermore, the integration of computational modeling could establish a platform for refining structural predictions and accelerating drug discovery efforts.

## Author Contributions

T.C. and J.L. designed the study; A.N., G.G.M.M., and N.W. prepared protein samples; S.S., A.V.B., C.K., and A.L. prepared gold-labeled samples; S.S., A.V.B., C.K., and M.S. performed ASAXS measurements; A.N. and G.G.M.M. performed FRET measurements; S.S., A.V.B., and G.G.M.M. analyzed the data; T.C. and J.L. supervised research; S.S., T.C., and J.L. wrote the manuscript with input from all authors.

## Acknowledgements

We thank Sebastian Doniach, Pehr Harbury, Rebecca Mathew and Theyencheri Narayanan for useful discussions, Thomas Zettl for initial work on protein labeling, and Thomas Nicolaus for laboratory assistance. We acknowledge the ESRF for the provision of synchrotron radiation facilities.

