## Supplementary Information for "Determination of Absolute Intramolecular Distances in Proteins by Anomalous X-ray Scattering Interferometry"

**CONTENT**

**Supplementary Materials and Methods**
**Supplementary Tables S1-S3**
**Supplementary Figures S1-S13**

### Supplementary Materials and Methods

**Overexpression, isolation and refolding of MalE proteins.**

Each single cysteine variant (MalE T31C, I212C, T36C, and S352C) and double variant (MalE T31C/I212C and MalE T36C/S352C) of MalE protein was expressed and purified according to published procedures (1). The T7/lac bacterial expression vector pET-23b(+) carrying a C-terminus 6x-His Tag sequence and containing the DNA coding sequence of MalE wild-type protein (2) was used as the template to generate the cysteine variants via PCR site-directed mutagenesis. *E. coli* BL21(DE3)pLysS transformant cells were grown at 37°C in LB medium containing ampicillin (0.1 mg/ml) and chloramphenicol (0.05 mg/ml). When an optical density (OD600) of 0.6-0.8 was reached, protein overexpression was induced by addition of 0.25 mM isopropyl β-D-1-thiogalactopyranoside (IPTG). The cells were harvested after 1.5 h and resuspended in 50 mL lysis buffer (50 mM Tris-HCl, pH 8.0, 1 M KCl, 10% glycerol, 10 mM imidazole and 1 mM dithiothreitol) for 2 L harvesting culture. Prior to cell disruption, cells were incubated for 30 minutes at 4°C in the presence of DNase I 500 ug/ml, cOmplete™, EDTA-free Protease Inhibitor Cocktail (1 tablet for each 50 mL bacterial culture), 0.2 mM phenylmethylsulfonyl fluoride (PMSF), and 2 mM dithiothreitol (DTT). Cells were lysed using an ultrasonic homogenizer (Branson Digital Sonifier; 25% amplitude, total exposure time to ultrasound of 10 minutes, with time lapses of 0.5 s ON/OFF pulse switches). Cell debris were discarded by a first step centrifugation at 5,000 × *g* for 30 min at 4°C. Next, the cell lysate supernatant was collected by a further ultracentrifugation step at 208,400 x *g* for 1 hour at 4°C. Each MalE protein was purified by Immobilized Metal Affinity Chromatography (IMAC) using the Ni Sepharose® 6 Fast Flow resin pre-washed with 10 column volumes lysis buffer. The cell lysate supernatant was loaded onto $Ni^{2+}$-sepharose resin (50 mL supernatant for 4 mL resin wet volume) and incubated overnight at 4°C. After two subsequent washing steps with 10 CV lysis buffer supplemented with 5 mM β-mercaptoethanol (BME) and 0.2 mM PMSF, the 6x-His Tagged MalE protein was eluted with 5 CV elution buffer (50 mM Tris-HCl, pH 8.0, 50 mM KCl, 10% glycerol, 250 mM imidazole, 0.2 mM PMSF and 5 mM BME). The eluted protein was dialyzed overnight at 4°C against 250 volumes of dialysis buffer (50 mM Tris-HCl, pH 8.0, 50 mM KCl and 5 mM DTT) to eliminate the excess of imidazole and using the SnakeSkin 10 kDa MWCO dialysis membrane tubing.

After the overnight dialysis at 4°C, the freshly purified MalE proteins were directly subjected to unfolding and refolding processes prior to storage, using adapted procedures based on Ganesh *et al.* (3). The isolated MalE variants were diluted down to 10-20 µM final concentration in unfolding buffer (6 M final concentration guanidine hydrochloride in 10 mM HEPES, pH 7.3). Each unfolding mixture was incubated for 3 hours under gentle rocking at 30°C. The unfolded protein solutions were afterwards centrifuged at 3046 *g* for 45 minutes at 4°C to remove irreversible aggregates that could hinder correct protein refolding. In parallel, a SnakeSkin 10 kDa MWCO dialysis membrane tubing was pre-soaked in water at 4°C for 1-2 min. The protein refolding was conducted at 4°C by a two-step dialysis process and over a total incubation period of 2 days. First, the proteins were each dialyzed against 100 volumes of refolding buffer (10 mM HEPES, 150 mM NaCl, pH 7.3) supplemented with 200 mM L-arginine. After 18-20 hours, a second dialysis step was carried out by exchanging the dialysis buffer solution with further 100 volume excess of refolding buffer. Both dialysis steps were conducted in the presence of 5 mM DTT. The refolded MalE proteins were collected after 42 hours and each concentrated to 500 µL final volume (starting from 2 L bacterial culture) using the Vivaspin centrifugal concentrator 10 kDa MWCO. The protein concentration was conducted at 3000 *g* at 4°C. The concentrated proteins were further loaded onto the Superdex™ 75 Increase 10/300 GL column using the automatized ÄKTA pure™ chromatography system (formerly GE Healthcare). For each protein, the column was pre-equilibrated with 2 CV storage buffer (labeling buffer I: 50 mM Tris-HCl, pH 7.4, 50 mM KCl supplemented with 2 mM DTT). Lastly, the MalE variants were concentrated to final concentration between 300 µM and 500 µM and finally stored at -80°C until further use. The final protein concentrations were determined with a NanoPhotometer® N60/N50 (IMPLEN), using the molar extinction coefficient of MalE (66350 $M^{-1} \cdot cm^{-1}$).

**FRET labelling and purification of MalE variants $MalE_{31\text{-}212}$ and $MalE_{36\text{-}352}$.**

Generally, we followed an already established protocol for stochastic maleimide labelling and purification of MalE proteins (1,2,4). $His_6$-tagged MalE double-cysteine variants were incubated in labelling buffer (50 mM Tris-HCl pH 7.4, 50 mM KCl) supplemented with 1 mM dithiothreitol (DTT) to retain the reduced state of both cysteine residues. Subsequently, the MalE variants were immobilized by nickel affinity on a nickel functionalized agarose medium ($Ni^{2+}$-Sepharose™ 6 Fast Flow, Cytiva). After that, the maleimide reaction with 50 nmol of each Alexa Fluor 555 (ThermoFisher) and Alexa Fluor 647 (ThermoFisher) was carried out in labelling buffer overnight at 4°C. The labeled, resin-bound proteins were washed with one column volume (void volume) labelling buffer and eluted with 500 µl labelling elution buffer (50 mM Tris-HCl pH 8.0, 50 mM KCl, 500 mM imidazole). Following the maleimide labelling with Alexa Fluor 555 and Alexa Fluor 647, the raw, fluorophore-labeled proteins were purified by size-exclusion chromatography (ÄKTA pure™ chromatography system, Cytiva; Superdex™ 75 Increase 10/300 GL, Cytiva) or anion exchange chromatography (ÄKTA pure™ chromatography system, Cytiva; MonoQ™ 5/50 GL column, Cytiva), respectively.

The eluate from the maleimide labelling protocol was prepared for further purification by removal of the remaining KCl and imidazole from the labelling elution buffer that could otherwise interfere with the anion exchange process. This step was done using a Sephadex G-25 medium (PD MiniTrap™ G-25, Cytiva) column. The labeled protein was then eluted in 1 ml of anion exchange sample buffer (10 mM Tris-HCl pH 7.5). The anion exchange column was set up with a 5 CV (resin volume) $ddH_2O$ wash, a 10 CV (resin volume) equilibration with anion exchange sample buffer, 10 CV (resin volume) equilibration with anion exchange elution buffer (10 mM Tris-HCl pH 7.5, 1 M NaCl) and a final 20 CV (resin volume) equilibration with anion exchange sample buffer. The washing step and all equilibrations were performed at 1 ml/min flow rate. Labeled protein was loaded onto the column with 0.5 ml/ min and the resin-bound protein was subsequently washed with 10 CV anion exchange sample buffer at 1 ml/min flow rate. For the consecutive elution a linear increase in anion exchange elution buffer ratio with a slope corresponding to 7.5 mM NaCl per column volume was chosen. The flow rate was adjusted to 0.5 ml/ min. Fractions containing the Alexa Fluor 555 and Alexa Fluor 647 labeled protein were selected and used for further analysis.

**smFRET measurements and data analysis**

Solution-based single-molecule FRET experiments were conducted on a homebuilt ALEX confocal microscope, as described in (5). All samples were measured in a 100 µL PBS droplet with a protein concentration ranging from 50 to 100 pM on a coverslip passivated with BSA (1 mg/ml in PBS). The experimental setup utilizes alternating laser excitation (ALEX) of two diode lasers: OBIS 532-100-LS (Coherent, USA), operated at 60 µW for donor molecules at 532 nm, and OBIS 640-100-LX (Coherent, USA), operated at 25 µW for acceptor molecules at 640 nm, both in alternation mode with a 100 µs total alternation period (20 kHz frequency). The lasers were combined by an aspheric fiber port (PAF2S-11A, Thorlabs, USA), coupled into a polarization-maintaining single-mode fiber P3-57 488PM-FC-2 (Thorlabs, USA), collimated (RC12APC-P01, Thorlabs, USA), before entering an epi-fluorescence confocal microscope (Olympus IX71, Hamburg, Germany), and guided into a water immersion objective (60X, NA 1.2, UPlanSAPO 60XO, Olympus, Japan) by a dual-edge beamsplitter ZT532/640rpc (Chroma/AHF, Germany). Fluorescence emitted from the sample was collected by the same objective and spatially filtered using a pinhole with a 50 µm diameter and spectrally split into the donor and acceptor channels by a single-edge dichroic mirror H643 LPXR (AHF, Germany). Fluorescence emission was filtered (donor: BrightLine HC 582/75 (Semrock/AHF, Germany); acceptor: Longpass 647 LP Edge Basic (Semrock/AHF, Germany) and focused onto avalanche photodiodes (SPCMAQRH-64, Excelitas, Canada). The detector outputs were recorded by an NI-Card PCI-6602 (National Instruments, USA) using LabView data acquisition software of the Weiss laboratory as outlined previously (6).
Data analysis was performed using a home-written software package, as described in (4). Bursts corresponding to single-molecule transits through the confocal volume were identified first using an all-photon burst-search (APBS) with a threshold of 15, a time window of 500 µs, and a minimum total photon number of 150 to determine donor leakage crosstalk (α) and direct acceptor excitation (δ). Then, a Dual-Channel-Burst-Search (DCBS) was performed with similar parameters to determine excitation flux (β) and detection efficiency and quantum yields differences (γ) to obtain accurate FRET values E as described (7) .
*E* histograms of double-labeled FRET species with Alexa Fluor 555 and Alexa Fluor 647 were extracted by selecting $0.3 < S < 0.7$ for 1D-E projections. *E* histograms of the open state without ligand (apo) and the closed state with 1 mM maltose (holo) were fitted with a Gaussian distribution $A \cdot e^{-\frac{(E-\mu)^2}{2\sigma^2}}$ . Distance conversion was done according to $R = R_0 \sqrt[6]{(1-E)/E}$, where R0 is the Förster radius (8) given by:

$$R_0{}^6 = \frac{9\, ln(10)}{128\pi^5\, N_A}\frac{\kappa^2}{n^4} Q_D \frac{\int_0^\infty F_D(\lambda)\varepsilon_A(\lambda)\lambda^4 \mathrm{d}\lambda}{\int_0^\infty F_D(\lambda)\mathrm{d}\lambda}$$

Here, NA is the Avogadro constant, κ2 the dipole orientation factor, n the averaged refractive index of the medium, QD the donor quantum yield, FD the donor emission spectrum, and the acceptor absorbance spectrum $\varepsilon_A$. The Förster radius for Alexa Fluor 555 – Alexa Fluor 647 was R0 = 51Å according to (1). The uncertainties in accurate FRET efficiencies were estimated through the error propagation of the γ factor, with a relative uncertainty of $\Delta\gamma/\gamma$ = 23% as described in (7).

**Synthesis of gold nanoparticles**

Gold nanoparticles were synthesized as previously described (9). Briefly, 0.54 g of gold(III) chloride hydrate and 1 g of β-D-thioglucose were dissolved in 72 ml of a 5:1 methanol:acetic acid solution (concentrations: 22 mM Gold(III) chloride and 63.5 mM β-D-thioglucose). The solution was stirred for 20 min at room temperature. Then, 0.9 g of sodium borohydride were dissolved in 20 ml distilled water (1.19 M sodium borohydride solution). The solution was added dropwise using a dropping funnel to the thioglucose-gold solution over 15 min with constant flow rate while stirring. The solution was stirred for another 30 minutes. Afterwards the solution (~90 ml) is concentrated to 10-15 ml using a rotary evaporator with 10 mbar at room temperature (~30-45 min). The rotation speed was set to 100 rotations per minute. Aggregates were filtered out using 0.45 µm CME filter units and the solution was desalted using an HPLC system with a Sephadex™ G25 column. The final glucose-coated gold nanoparticles are dissolved in ddH$_2$O and are stable for weeks at 4 °C or for months at −20°C.

**Protein–gold nanoparticle conjugation and purification**

Maltose binding proteins (MBPs) were conjugated to the gold nanoparticles synthesized as described in the previous section. The His-tags of the proteins were immobilized on a fast flow nickel sepharose column. Binding the proteins on nickel sepharose resin during labeling, reduces aggregation and improves the labeling efficiency. The column, containing the protein (protein solution: 500 µl with 100 µM), was flushed with 50 mM Tris-KCl buffer, pH 8. The gold nanoparticles were buffer exchanged to the same buffer with Amicon® centrifugal filters (3kDa MWCO) and introduced to the column (500 µl with 2000 µM). Importantly, the molar concentration of gold was at least 20-fold higher than the protein concentration to improve the labeling efficiency and reducing aggregation. The column was sealed and left undisturbed at 4 °C over night. The excess gold nanoparticles were flushed out with 50 mM Tris-KCl buffer and reused. The labeled proteins were eluted from the column and collected with 50 mM Tris-KCl buffer supplemented with 250 mM imidazole. The collected proteins were subsequently purified using an HPLC system with a Superdex™ 75 column and we selected the fraction with the highest labeling efficiency by the A260/A280 ratio and absorption at 360 nm (procedure and chosen fractions are shown in Supplementary Figure S2 and S3). After purification, the selected fractions were concentrated with Amicon® centrifugal filters with a 3 kDa MWCO.

**ASAXS measurements**

ASAXS measurements were performed using a flow-through capillary cell with a diameter of 2 mm at the beamline ID02 of the European Synchrotron Radiation Facility, Grenoble (France) using a sample to detector distance of 1,2 m (10). 2D scattering images were recorded using an Eiger2X 4M detector at 9 energies (11619, 11819, 11869, 11899, 11909, 11914, 11919, 11929, 11969 eV) around the L-III absorption edge of gold to over-determine the matrix equation (see below). 2D data were reduced to azimuthally averaged 1D scattering profiles using standard techniques. To achieve highest accuracy, the normalization to absolute units in [mm$^{-1}$] was repeated at each energy and for each sample using a second water-filled flow-through capillary (11-13). The gold labeled protein samples with a concentration of ~200 nM were loaded into the capillary using a syringe pump and moved during the experiment to avoid radiation damage. After ramping the X-ray energy over the absorption edge, the measurement at 11619 eV was repeated and compared with the first recording to exclude a deterioration of the sample during the ASAXS measurements.

Matching buffer solutions for background correction were measured using the same flow-through cell after concentration with amicon filters at the same X-ray energies. In this way, the buffer scattering could be subtracted from the sample scattering with highest accuracy before further analysis.

**ASAXS data analysis**

The atomic scattering factor close to an atomic absorption edge is well described by:

$$f(E) = f_0 + f'(E - E_0) + if''(E - E_0)$$

Away from an absorption edge the f' and f'' terms are negligible and the atomic scattering factor is given by $f_0$. At an absorption edge the energy of the incident photon is equal to an electronic transition. For gold at 11.919 keV one electron from the L-III orbit ($2p_{3/2}$ -electron) is ejected from the atom and the photon is absorbed. Therefore, close to the absorption edge, the *f'* and *f''* terms increase and the scattering factor becomes anomalous.

To perform ASAXS with gold labels we measure at 9 different energies around the L-III absorption edge. At the edge the fraction of absorbed photons by gold is larger and less photons are scattered than off-edge. The scattering from other atoms remains approximately unchanged. Here we use a matrix inversion approach (14,15) to isolate the partial structure factor of the gold-gold interference from other scattering contributions. A maximum entropy algorithm is employed to Fourier transform the gold-gold interference into a real-space distance distribution.

The scattering profiles of the 9 selected energies are normalized as described in the supplementary information of (14) using the theoretical intensity scaling factors for gold given by:

$$int(E) = |f_{Au}(E)|^2 + 2Re(f_{Au}(E))$$

ASAXS data reduction was carried out similar to Pinfield (15) and Zettl (14). Briefly, we determined the gold-gold scattering contribution from the corrected and energy-dependent scattering data with a matrix inversion approach based on the following matrix relation:

$$\begin{pmatrix} I_{E_1}(q) \\ I_{E_2}(q) \\ \vdots \\ I_{E_{N_E}}(q) \end{pmatrix} = \begin{pmatrix} a_{E_1} & b_{E_1} & c_{E_1} \\ a_{E_2} & b_{E_2} & c_{E_2} \\ \vdots & \vdots & \vdots \\ a_{E_{N_E}} & b_{E_{N_E}} & c_{E_{N_E}} \end{pmatrix} \times \begin{pmatrix} G_{Au-Au}(q) \\ G_{Au-prot}(q) \\ G_{prot-prot}(q) \end{pmatrix}$$

$$I = T \times G$$

Here the scattering intensity vector I has the scattering profiles for all energies concatenated with dimensions $(N_E \times N_q, 1)$, where $N_E$ is the number of energies and $N_q$ is the number of measured q channels. The transformation matrix T consists of the diagonal submatrices a, b and c with dimensions $(N_q, N_q)$, where

$$a_{E_i} = |f_{Au}(E_i)|^2 \cdot F_a(q)$$

with $f_{Au}$ being the atomic scattering factor of gold and $F_a$ is the squared form factor of the gold spheres calculated with the measured radius distribution of the gold labels. Similarly, we have:

$$b_{E_i} = f_{Au}(E_i) \cdot f_{prot} \cdot F_b(q)$$

and

$$c_{E_i} = \left|f_{prot}\right|^2 \cdot F_c$$

With the atomic scattering factor of the protein:

$$f_{prot} = \frac{1}{N_{atoms}} \cdot (N_H \cdot f_H + N_C \cdot f_C + N_N \cdot f_N + N_O \cdot f_O + N_S \cdot f_S)$$
$$= \frac{1}{5742}(1852 \cdot f_H + 2865 \cdot f_C + 469 \cdot f_N + 548 \cdot f_O + 8 \cdot f_S)$$

where the atomic composition of the MBP mutant $MalE_{36\text{-}352}$ was used. $F_b$ is the form factor of the gold spheres and $F_c$ is a unit matrix since the protein form factor is neglected. Scattering factor data is taken from:
http://skuld.bmsc.washington.edu/scatter/AS_periodic.html

Additionally, we can disregard the energy dependence of the protein scattering factor in our case, as atomic scattering factors for non-gold atoms in the sample show minimal energy dependence within the chosen energy range (Supplementary Figure S7). Lastly, the G vector $(3 \times N_q, 1)$ contains partial structure factors of gold-gold ($G_{Au-Au}$), gold-protein ($G_{Au-prot}$) and protein-protein ($G_{prot-prot}$). The matrix I is experimentally determined from ASAXS measurements of double-labeled samples, while the transfer matrix T is computed using tabulated atomic scattering factors and treating the gold labels as spheres with a radius distribution around 7 Å (Supplementary Figure S1). Matrix inversion then yields the matrix G, which contains the gold-gold structure factor. The q-range was cut to 0.077 $Å^{-1}$ to 0.315 $Å^{-1}$ to focus on the oscillation. The gold-gold structure factor $G_{Au-Au}$ was corrected by a constant offset by subtracting the mean and Fourier transformed with a maximum entropy algorithm to obtain the pair distance distributions P(d) .

**Subtraction of unlabeled and single-labeled proteins**

Achieving high sample purity proves challenging for our gold-labeled proteins. The double-labeled sample also contains single-labeled and unlabeled proteins even after purification. To account for the presence of single-labeled and unlabeled proteins, we subtracted different amounts of single-labeled and unlabeled profiles from the scattering curves of the double-labeled sample. Protein gels give a first estimate for the ratio of double-labeled to single-labeled to unlabeled proteins as 10% to 30% to 60% (Figure S2). We chose single-labeled mutants MalE352 and MalE31 for the subtraction. Importantly, we note that subtracting different amounts of unlabeled or single-labeled protein is not shifting the main peak position for more than 1 Å. (Figure S10, S11 and Table S1, S2). We find that subtracting 60% of unlabeled and 30% of single-labeled sample minimizes secondary peaks. We used Guinier analysis (16) to scale AXSI measurements of single-labeled sample and unlabeled sample to match the concentration of the double-labeled sample, where we used the following values:

$$\rho_{solvent} = 0.334\ \frac{e}{\text{Å}^3}$$

$$\rho_{gold} = 4.128\ \frac{e}{\text{Å}^3}$$

$$\rho_{prot} = 0.422\frac{e}{\text{Å}^3}$$

where $\rho$ is the electron density of solvent, gold and the protein. $\rho_{gold}$ was calculated from the scattering length density at 11619 eV using an online calculator http://www.refcalc.appspot.com/sld. The electron densities for protein and solvent were taken from CRYSOL (17). Furthermore, we also needed the volumes of the protein and the gold-spheres:

$$V_{gold} = 1437\ \text{Å}^3$$

$$V_{prot} = 59.230\ \text{Å}^3$$

For the double-labeled sample, the forward scattering is proportional to:

$$I(0) \sim (\Delta\rho_{prot} \cdot V_{prot} + 2 \cdot \Delta\rho_{gold} \cdot V_{gold}\ )^2.$$

For single-labeled proteins:

$$I(0) \sim (\Delta\rho_{prot} \cdot V_{prot} + \Delta\rho_{gold} \cdot V_{gold}\ )^2.$$

And for unlabeled proteins:

$$I(0) \sim (\Delta\rho_{prot} \cdot V_{prot})^2.$$

Combined, we obtain the following $I(0)$ ratios for monodisperse samples with identical concentration:

$$I(0)_{unlabeled} = 0.105\ I(0)_{double}$$

$$I(0)_{single} = 0.44\ I(0)_{double}$$

Additionally, we computed the forwardscattering from pdb files that contained MalE and modeled gold nanocrystals with CRYSOL (17), which improves the calculation by taking the solvation shell into account.

$$I(0)_{unlabeled} = 0.12\ I(0)_{double}$$

$$I(0)_{single} = 0.45\ I(0)_{double}$$

The gold nanoparticles consist of approx. 85 atoms (size measurements, Figure S1). Atom number can be calculated with the following formula (with *D* being the diameter of the particle):

$$N = 30.92\,(D[nm])^3$$

**Coarse-grained simulation of accessible volumes**

For comparison to FPS and FRETraj distances we computed the accessible volumes also with a simple “sterics only” simulation custom written in python. The code reads in all atom positions and then creates a grid of possible label positions around the two attachment points, taken to be the $C_\beta$ atoms of the cysteines introduced for labeling. The grid spacing is 1 Å. All positions that are further away from the attachment atoms than the linker length are deleted. Furthermore, all positions are deleted that are closer than the gold sphere radius (7 Å) to any atom in the protein. The remaining points form the accessible volume.

#### Supplementary Tables

| | Subtraction: unlabeled single-labeled [%] | Distance, mean [Å] | Distance, uncertainty [Å] | Peak width, sigma [Å] | Peak width, uncertainty [Å] |
|---|---|---|---|---|---|
| **$MalE_{36-352}$** | **60-30** | 56.2 | 0.3 | 1.9 | 0.1 |
| | **30-20** | 56.0 | 0.2 | 2.1 | 0.1 |
| | **0-0** | 55.8 | 0.2 | 2.2 | 0.1 |
| **$MalE_{36-352}$ repeat** | **60-30** | 56.2 | 0.2 | 2.0 | 0.2 |
| | **30-20** | 55.8 | 0.3 | 2.1 | 0.1 |
| | **0-0** | 55.9 | 0.1 | 2.1 | 0.1 |
| **$MalE_{36-352}$ result** | | **56.0** | **0.3** | **2.0** | **0.1** |
| **$MalE_{36-352}$ + 10 mM maltose** | **60-30** | 43.3 | 0.7 | 3.3 | 1.0 |
| | **30-20** | 43.5 | 0.4 | 5.0 | 0.9 |
| | **0-0** | 42.7 | 0.6 | 6.6 | 1.5 |
| **$MalE_{36-352}$ + 10 mM maltose repeat** | **60-30** | 43.2 | 0.6 | 3.5 | 0.8 |
| | **30-20** | 43.1 | 0.5 | 4.9 | 0.9 |
| | **0 - 0** | 42.8 | 0.4 | 5.5 | 0.6 |
| **$MalE_{36-352}$ + 10 mM maltose result** | | **43.1** | **0.6** | **4.8** | **1.5** |

**Supplementary Table S1. Measured AXSI distances for $MalE_{36-352}$.** Distances are computed by fitting a Gaussian to the main peak of the *P*(d) distributions as illustrated in Figure S9 for 20 individual maximum entropy runs. The reported mean and uncertainty are the mean and standard deviation of 20 main peak positions from repeat runs of the maximum entropy code. Similarly, the width of the Gaussian peaks sigma is given with uncertainty. Both mutants with and without maltose were corrected for 60 % unlabeled and 30 % single-labeled, 30 % unlabeled and 20 % single-labeled or not corrected at all. The resulting distances are shown in the table. The result column shows the average for all subtractions and repeats. Values are discussed in the main text.

| | Subtraction: unlabeled-single-labeled [%] | Distance, mean [Å] | Distance, uncertainty [Å] | Peak width, sigma [Å] | Peak width, uncertainty [Å] |
|---|---|---|---|---|---|
| **MalE$_{31\text{-}212}$** | **60-30** | 63.3 | 0.2 | 2.9 | 0.2 |
| | **30-20** | 64.1 | 0.2 | 1.9 | 0.1 |
| | **0-0** | 63.7 | 0.6 | 1.9 | 0.1 |
| **MalE$_{31\text{-}212}$ result** | | **63.7** | **0.5** | **2.2** | **0.5** |
| **MalE$_{31\text{-}212}$ + 10 mM maltose** | **60-30** | 59.3 | 0.4 | 2.8 | 0.4 |
| | **30-20** | 59.4 | 0.3 | 1.7 | 0.1 |
| | **0-0** | 59.1 | 0.2 | 1.8 | 0.1 |
| **MalE$_{31\text{-}212}$ + 10 mM maltose repeat** | **60-30** | 60.0 | 0.4 | 2.5 | 0.7 |
| | **30-20** | 59.3 | 0.2 | 1.7 | 0.1 |
| | **0–0** | 59.0 | 0.4 | 1.8 | 0.1 |
| **MalE$_{31\text{-}212}$ + 10 mM maltose result** | | **59.4** | **0.5** | **2.0** | **0.6** |

**Supplementary Table S2. Measured AXSI distances for MalE$_{31\text{-}212}$.** Equivalent to Table S1 but for MalE$_{31\text{-}212}$.

| | Linker length [Å] | Width [Å] | Radius 1 [Å] | Radius 2 [Å] | Radius 3 [Å] |
|---|---|---|---|---|---|
| **Gold NP** | **10** | **2** | **7** | **7** | **7** |
| Alexa 555 – C2 Maleimide | **21** | **4.5** | **8.8** | **4.2** | **1.5** |
| Alexa Fluor 647 – C2 Maleimide | **21** | **4.5** | **11** | **4.7** | **1.5** |

**Supplementary Table S3. Geometrical Parameters for the linkers used to model FRET and AXSI measurements.** Values for Alexa fluorophores taken from (18).

#### Supplementary Figures

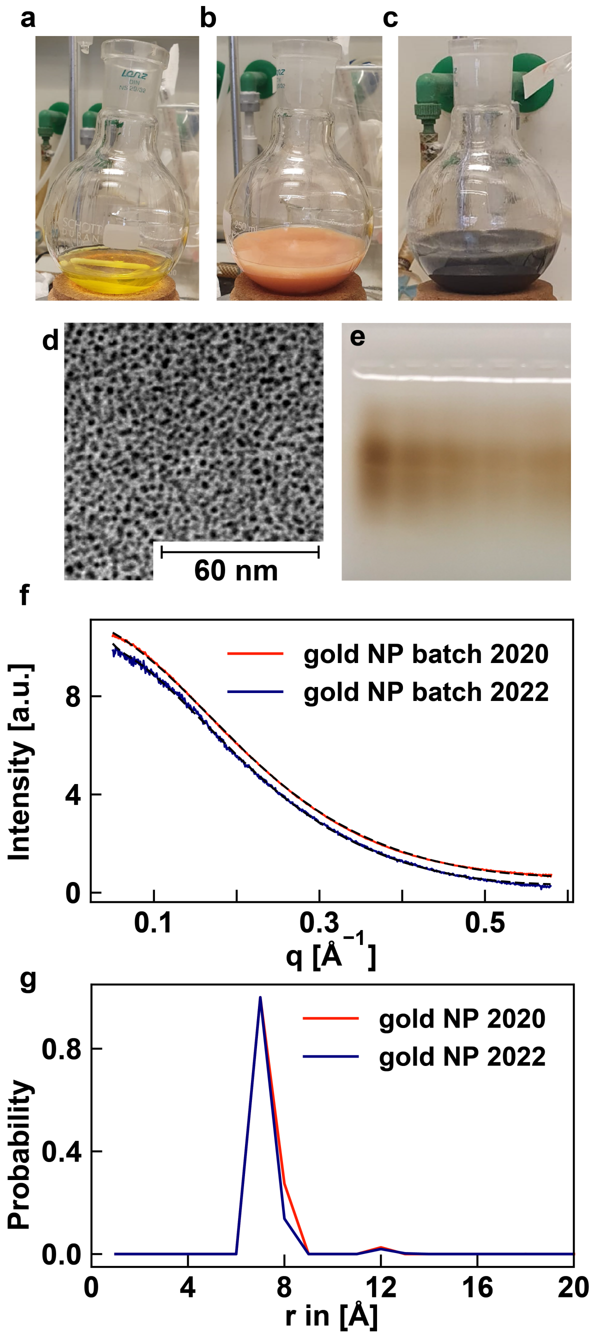


**Suppplementary Figure S1. Preparation and characterization of gold nanoparticles for protein labeling.** Gold solution during synthesis: **a)** Gold(III) chloride hydrate dissolved in 5:1 methanol:acetic acid. **b)** Gold(III) chloride hydrate solution after mixing with dissolved thioglucose. **c)** Same solution after dropwise addition of sodium borohydride. **d)** TEM image of gold nanocrystals. **e)** 1 % Agarose gel of different concentrations of gold nanocrystals. The gel was run in TAE buffer for 15 min at 70 mA. **f)** SAXS profiles for two independently synthesized batches of gold nanoparticles. SAXS data are shown in blue and red; black lines are fits of the profiles as a superposition of spheres with different radii. Data are vertically offset for clarity. **g)** Fitted distributions of sphere radii. The gold nanoparticles show narrow radius distributions with a mean of 7 Å and good reproducibility between batches.

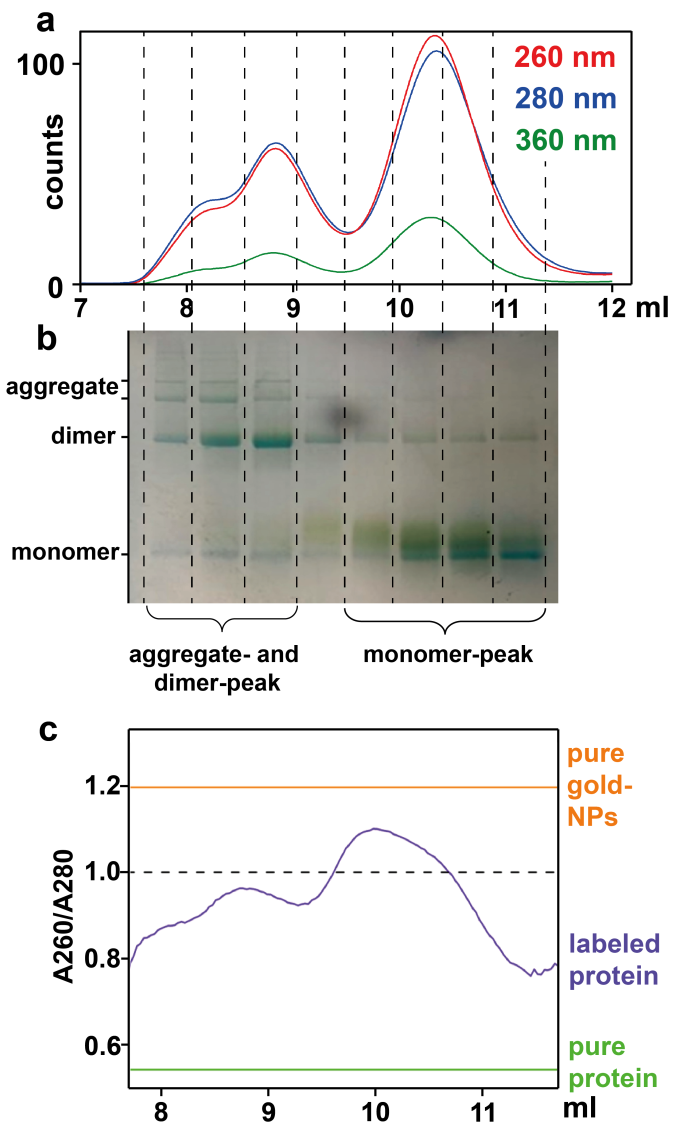


**Supplementary Figure S2. Protein labeling and purification.** **a)** HPLC elution profile with a Superdex® Increase 75 column and **b)** SDS page gel (Mini-Protean TGX® PAGE gel any kD, Bio-Rad®) electrophoresis (denatured) of gold-labeled MalE36-352. The first peak with a shoulder on the left from 7 to 9 ml corresponds to mostly aggregates and dimers with a lower fraction of gold. The right peak (between 9.5 and 11 ml) consists of mostly monomers. The gel was stained with Coomassie blue and LI Silver enhancement (greenish), which stains the gold NPs. The lanes in the gel correspond to fractions in the HPLC run (indicated by dashed lines). The upper two bands correspond to protein cluster with MW > 113 kDa. The third band corresponds to dimers with MW of 81-83 kDa. The lower triple band corresponds to protein monomers with MW from 41-53 kDa. We note, that molecular weight estimates are not accurate for gold NPs. **c)** Ratio of 260 nm to 280 nm absorbance. The ratio is above 1 for the left flank of the monomer peak. For ASAXS measurements only the left flank of the monomer-peak was selected. From the HPLC run we can also compute the protein:gold concentration ratio in the selected fractions. For the run shown here we get a ratio of 1:0.75 (see Figure S3 caption for calculation). However, small gold and protein concentrations were used. Labeling high concentrations of proteins, which is necessary for SAXS measurements, reduces the protein:gold ratio to ~1:0.5 (high concentration HPLC runs in Supplementary Figure S3).

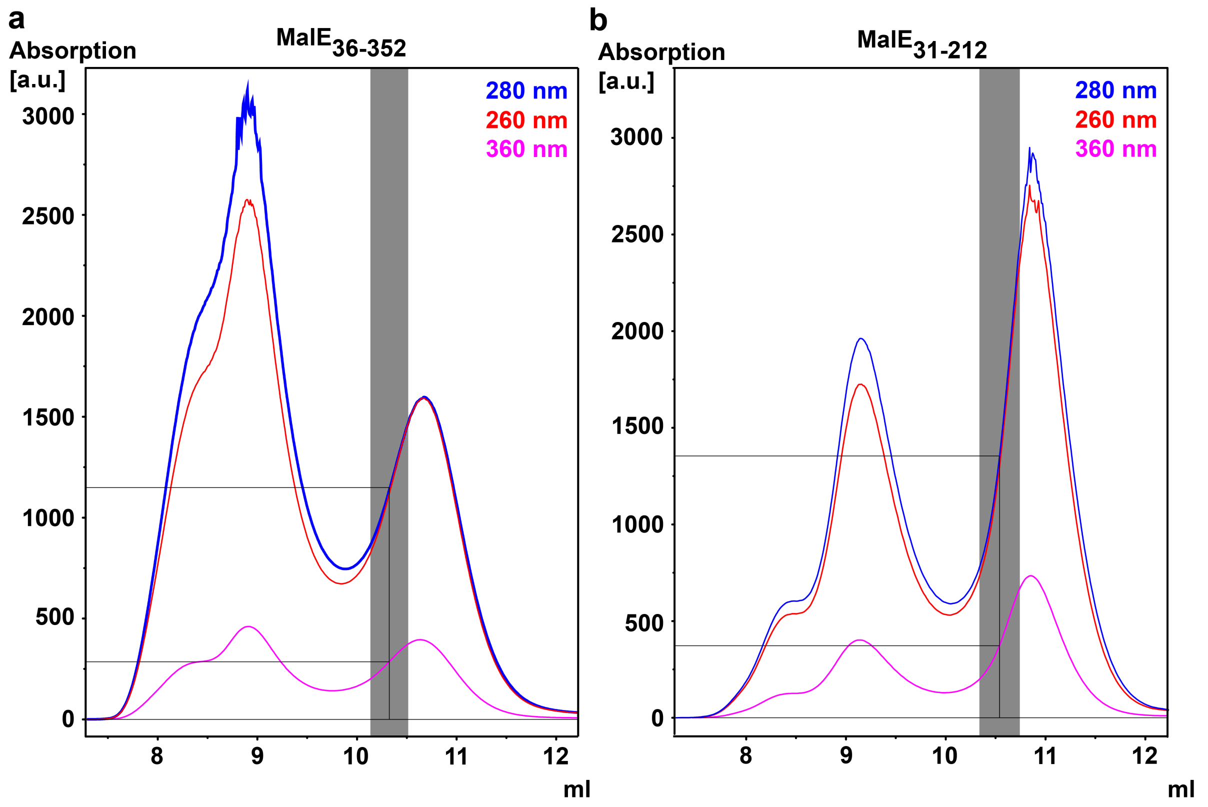


**Figure S3. Size exclusion chromatography runs for larger amounts of MalE$_{31\text{-}212}$ and MalE$_{36\text{-}352}$.** Absorption was measured at 260, 280 and 360 nm. Grey backgrounds indicate the fractions that were used for AXSI measurements. The extinction coefficient of gold NPs is $\varepsilon = 7.6 \cdot 10^4 \frac{1}{Mol \cdot cm}$ at 360 nm (9). The gold absorption at 280 nm is $\varepsilon = 15.85 \cdot 10^4 \frac{1}{Mol \cdot cm}$ (9) and the proteins used here do not absorb at 360 nm and have an extinction coefficient of $\varepsilon = 6.65 \cdot 10^4 \frac{1}{Mol \cdot cm}$ at 280 nm (computed with (19)). To compute the protein:gold ratio we used the counts in the center of the selected fractions (black lines). At 360 nm all counts originate from gold NPs. We can calculate the absorption that originates from gold NPs at 280 nm with the following formula: $counts_{gold,280\,nm} = counts_{gold,360\,nm} \cdot \frac{\varepsilon_{gold,280\,nm}}{\varepsilon_{gold,360\,nm}}$. The protein absorption at 280 nm is then given by: $counts_{prot,280\,nm} = counts_{total,280\,nm} - counts_{gold,280\,nm}$. Concentrations are given by the counts divided by the extinction coefficients. For MalE$_{36\text{-}352}$ we obtain a protein:gold ratio of 1:0.45 and for MalE$_{31\text{-}212}$ we obtain 1:0.56. We note that the protein:gold ratio for a perfectly double-labeled sample is 1:2. The measured ratios ~1:0.5 agree with our estimate of 30% single-labeled and 10% double-labeled sample from the scattering profiles.

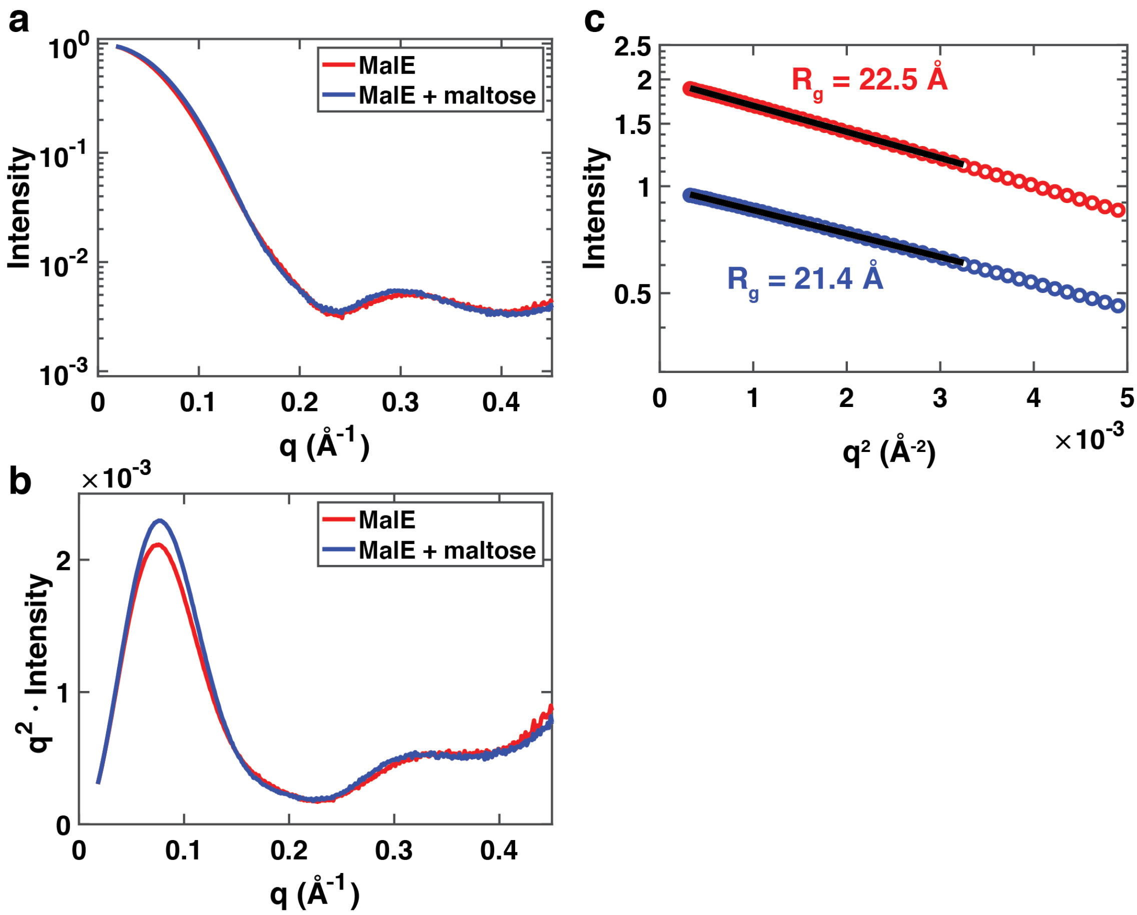


**Supplementary Figure S4. SAXS analysis for unlabeled MalE in the apo and holo configuration.** Normalized SAXS data for $MalE_{36\text{-}352}$ without gold labels at a fixed energy of 11.619 keV. **a)** Scattering profiles of $MalE_{36\text{-}352}$ in the absence of maltose (red) and in the presence of 10 mM maltose (blue). The profiles show subtle difference, but are overall similar. **b)** Kratky representation (scattering intensity weighted by $q^2$ vs. $q$) of the same data as shown panel a. The Kratky representation highlights the conformational change upon addition of maltose, as has been observed for similar bi-lobed proteins previously (20,21). **c)** Guinier analysis of the scattering profiles shown in panel a. The logarithm of the scattering intensity vs. $q^2$ for the lowest $q$ values is well described by a straight line (shown in black), indicative of a dilute and monodisperse sample. From the slope of the linear fits, we have determined the radius of gyration to be $R_g$ = 22.5 Å ± 0.3 Å in the absence of maltose and $R_g$ = 21.4 Å ± 0.3 Å in the presence of 10 mM maltose. Errors are from the uncertainty of the fits. The data set in the absence of maltose (red) is scaled by a factor of 2 for clarity. The experimentally determined radii of gyration are in excellent agreement with previous experimental results (20) and with the values computed from the crystal structures of the open (22) (PDB accession code 1OMP; Rg = 22.8 Å; radii of gyration were computed using CRYSOL (17) with default settings) and closed, maltose bound conformation (23) of MalE (PDB accession code 1ANF; Rg = 21.8 Å).

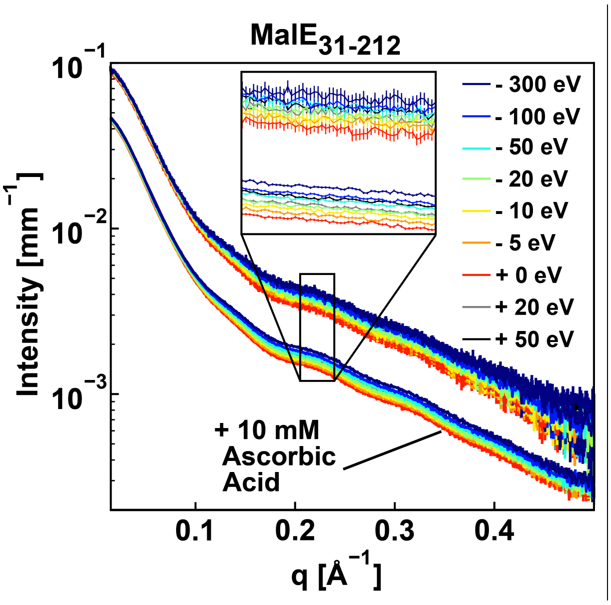


**Supplementary Figure S5. Effect of exposure time and averaging on the signal-to-noise ratio.** ASAXS data for double-labeled $MalE_{31\text{-}212}$. Upper curves are the average of 5 frames with 0.3 s exposure time each and the photon flux was $2.3\cdot10^{11}$ photons/s. The lower curves are the average of 8 frames with 0.5 s each and a flux of $7.3\cdot10^{12}$ photons/s. The higher flux and longer exposure time are made possible by addition of 10 mM ascorbic acid. The sample in the capillary was moved between frames. Radiation damage was not observed.

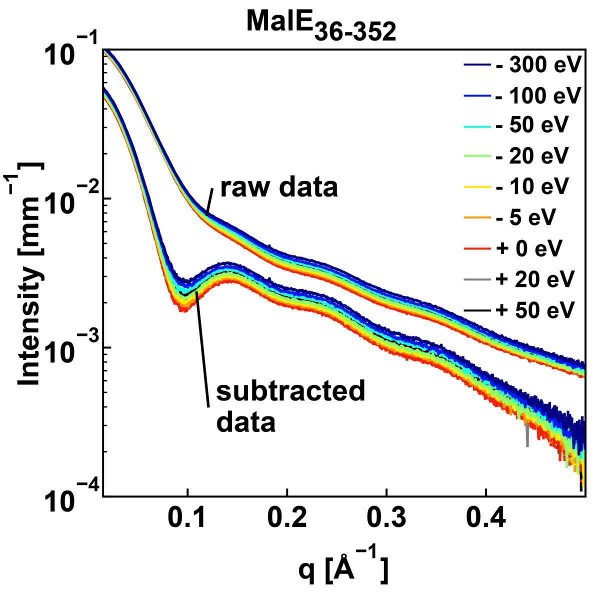


**Supplementary Figure S6. Scattering Data after the subtraction of unlabeled and single-labeled proteins.** Similar to the buffer subtraction we subtracted 60 percent of unlabeled and 30 percent of single-labeled MalE proteins from the measurements that include the double-labeled sample. Matching single-labeled and unlabeled proteins were also measured at the same energies.

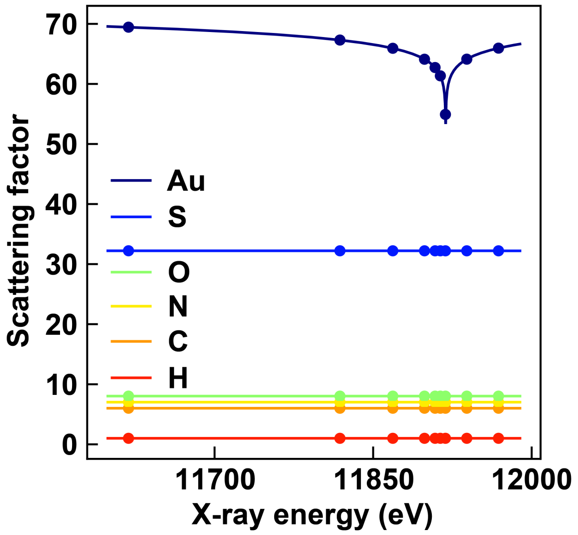


**Supplementary Figure S7. Real part of atomic scattering factors.** Atomic scattering factors for all atoms present in the sample. Points depict the 9 energies selected for AXSI measurements. The scattering factors for the atoms present in the protein show almost no energy dependence in the chosen energy regime, while the gold LIII-absorption edge of gold is clearly visible as a minimum at 11919 eV.

Data taken from: http://skuld.bmsc.washington.edu/scatter/AS_periodic.html

.

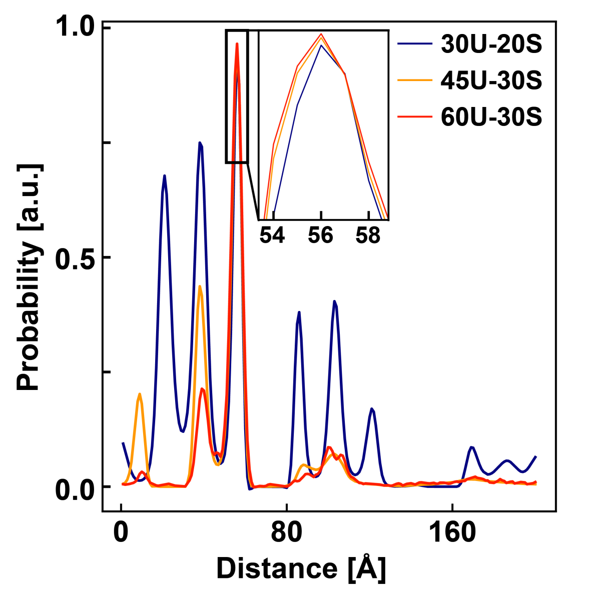


**Supplementary Figure S8. Distance distributions for MalE$_{36\text{-}352}$ for different background subtraction schemes.** Example distributions for different subtraction of unlabeled and single-labeled. 30 % unlabeled 20 % single-labeled (blue), 45 % unlabeled 30 % single-labeled (orange) and 60 % unlabeled 30 % single-labeled (red). The zoom in depicts the main peak positions.

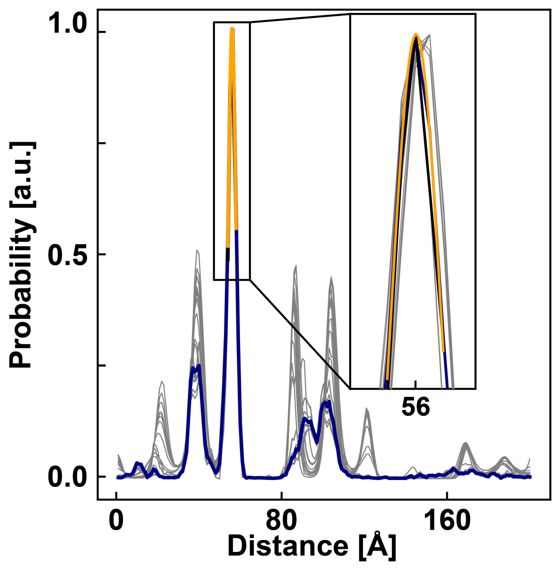


**Supplementary Figure S9. Distance distributions for MalE$_{36\text{-}352}$ from repeat runs of the maximum entropy inversion algorithm.** 20 distance distributions obtained from repeat runs of the maximum entropy inversion scheme using the scattering data after subtraction of 60% unlabeled and 30% double-labeled are shown (grey thin lines). The mean peak position is given in Table S1. One distribution is emphasized in blue. The Gaussian fit that is used to determine the peak center and standard deviation is shown in orange. The inset shows a zoom on the main peak position.

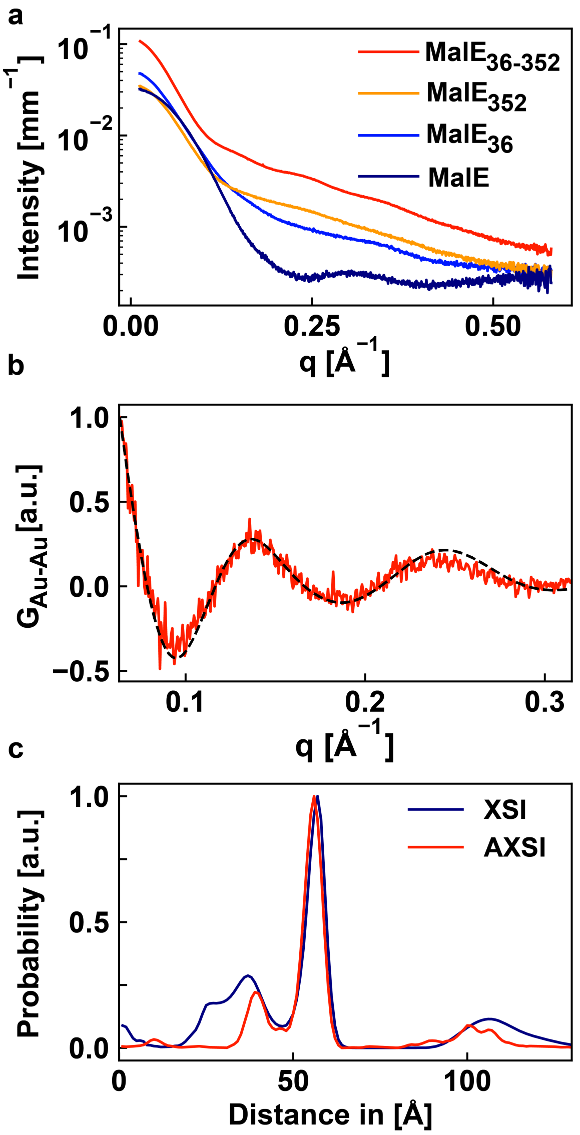


**Supplementary Figure S10. Distance distribution for MalE36-352 by single energy XSI.** a) SAXS measurements on MalE36-352 with two labels ($I_{double}$), one label MalE36 ($I_{single,1}$) at amino acid position 36, one label MalE352 at 352 ($I_{single,2}$) and unlabeled MalE ($I_{unlabeled}$) measured at 11.619 keV. b) Interference term $G_{Au-Au} = I_{double} - \left(I_{single,1} + I_{single,2} - I_{unlabeled}\right)$ calculated with the regular XSI method (red line) and the fit (black dashed line). Details of the regular XSI procedure in (24) c) Comparison of the distance distributions computed with AXSI for MalE$_{36\text{-}352}$, where we subtracted 60% single-labeled and 30% unlabeled protein, and the single energy XSI method.

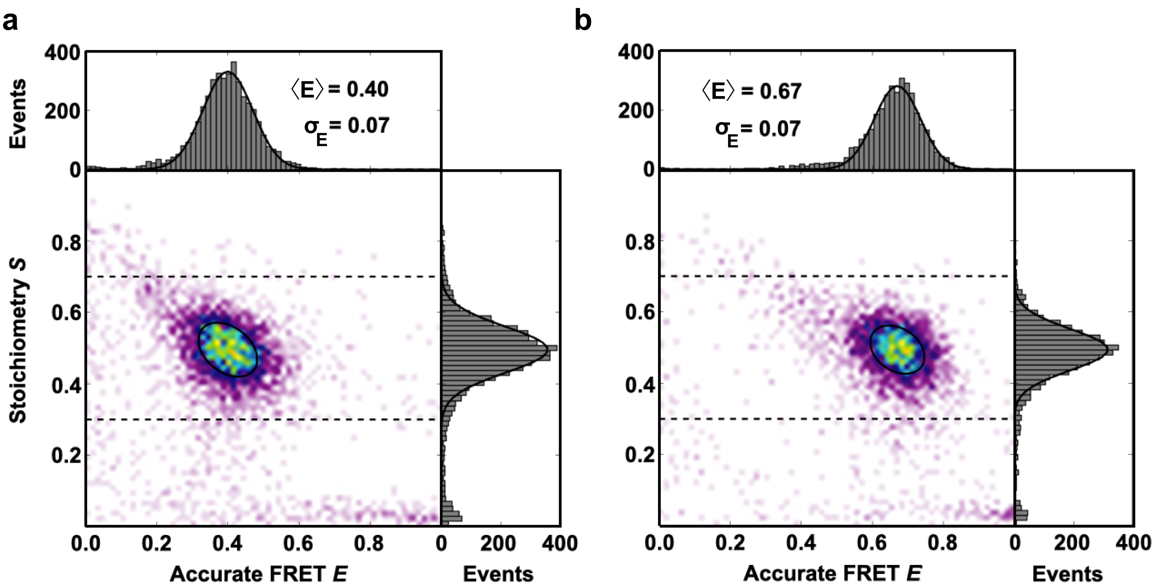


**Supplementary Figure S11. Monitoring conformational changes in MalE36-352 by smFRET/ALEX**. Corrected accurate *ES*-histograms of $MalE_{36\text{-}352}$ labeled with Alexa555 and Alexa647 for the open conformation in the absence of maltose (a) and the liganded, closed (b) conformation in the presence of 1 mM maltose.

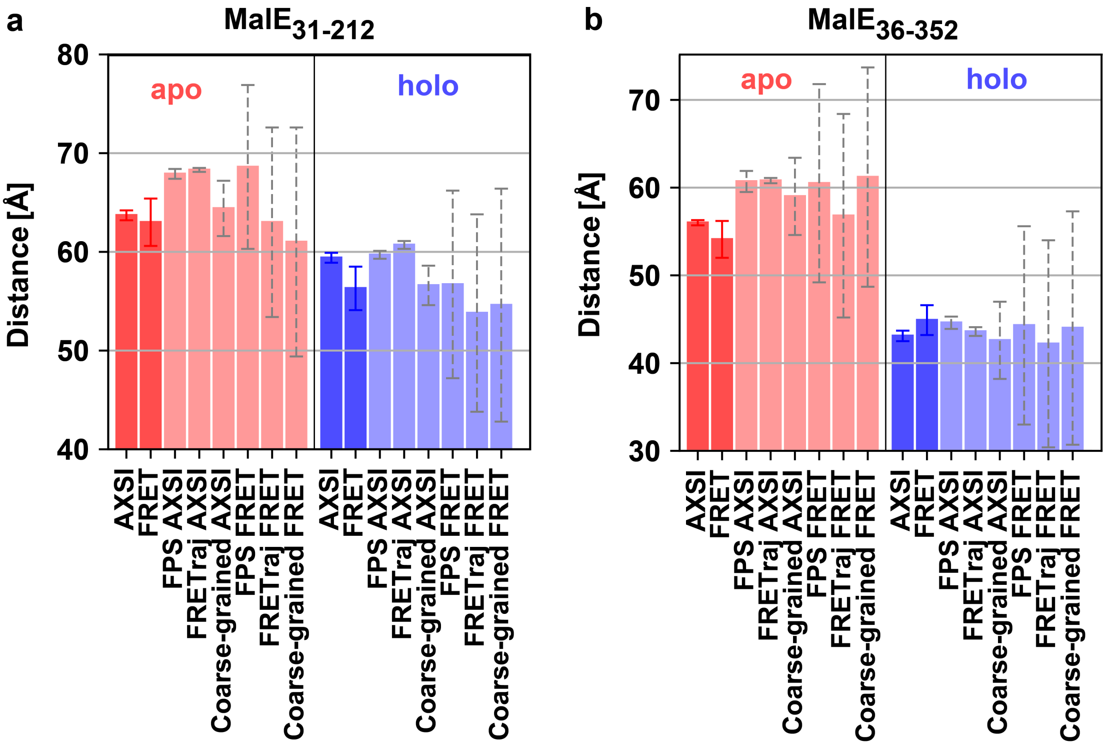


**Figure S12. Visualization of measured and computed distances between AXSI and FRET labels for different MalE constructs.** The open/apo state is depicted in red and the closed/holo state in blue. AXSI and FRET data shown in darker, simulation data in lighter blue/red. Error bars for the experiment data are experimental errors of the mean. Simulation errorbars depict the standard deviations of all possible distances between the accessible volumes (AVs) or accessible contact volumes (ACVs).

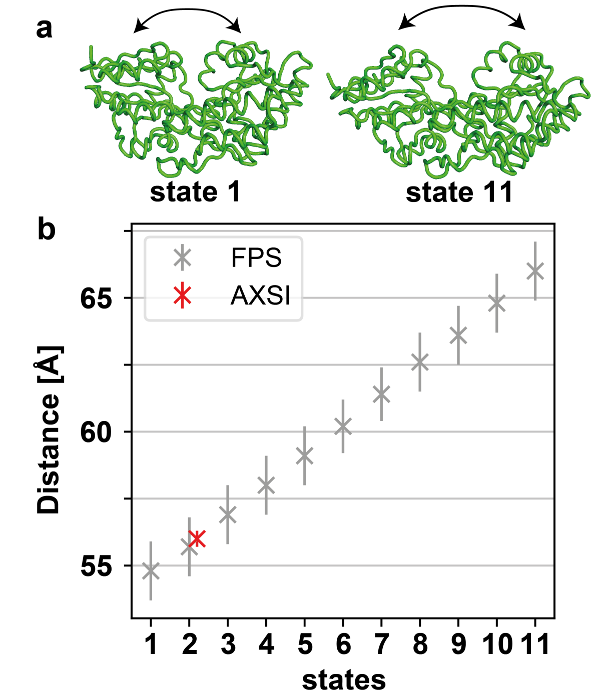


**Supplementary Figure S13. Elastic network simulation for MalE$_{36-352}$ in the apo state. a)** illustration of protein movement computed by elNemo (25) along the first non-trivial normal mode (mode 7). (PDB ID1OMP). B) FPS distances for a 10 Å long linker and 7 Å radius gold NPs for 11 states or conformations along the first non-trivial normal mode. In red the AXSI measurement for MalE$_{36-352}$ is shown.
